# Generative Modeling of Single Cell Gene Expression for Dose-Dependent Chemical Perturbations

**DOI:** 10.1101/2022.10.05.510890

**Authors:** Omar Kana, Rance Nault, David Filipovic, Daniel Marri, Tim Zacharewski, Sudin Bhattacharya

**Author notes:** **Corresponding author**: Sudin Bhattacharya, Michigan State University, 775 Woodlot Dr, Rm 1313, East Lansing, MI 48824, USA.

## Abstract

Single cell sequencing provides a new opportunity to study the heterogeneity of chemical perturbation within tissues. However, exploring the combinatorial space of all cell type-chemical combinations is experimentally and financially unfeasible. This space is significantly expanded by the dose axis of chemical perturbation. Thus, computational tools are needed to predict responses not only across tissues, but also across doses while capturing the nuances of cell type specific gene expression. Variational autoencoders simplify the single cell expression space allowing cross cell type predictions using simple vector arithmetic. However, differing sensitivities and non-linearities make cell type specific gene expression predictions following treatment at higher doses challenging. Here we introduce **single cell Variational Inference** of **Dose-Response** (*scVIDR*) which achieves high dose and cell type specific predictions better than other state of the art algorithms. scVIDR predicts in vivo and in vitro dose-dependent gene expression across cell types in mouse liver, peripheral blood mononuclear cells, and cancer cell lines. We use regression to interpret the outputs of scVIDR. Additionally, we use scVIDR to order individual cells based on their sensitivities to a particular chemical by assigning a pseudo-dose value to each cell. Taken together, we show that scVIDR can effectively predict the dose and cell state dependent changes associated with chemical perturbations.

## Introduction

In 2010, Sydney Brenner suggested that it is possible to deduce the physiology of biological systems by understanding the interactions and behaviors of their constituent units^1^. The appropriate unit, in his opinion, was the cell. Single cell sequencing (scSeq) has revolutionized the study of cell biology. With the ability to capture the transcriptomic state of thousands of cells at once, a fine-grained picture of the organization of cell physiology has begun to emerge^2^. Much of the effort in scSeq has been made in the realm of cell type/state discovery^3,4^, cellular development^5–8^, and disease progression^9,10^. These represent natural applications of scSeq, especially regarding the spatial and temporal dynamics of cellular systems and their interactions. However, relatively little attention has been given to how cells respond to environmental signals like chemical exposures, which in addition to being spatial and temporal, are also chemical- and dose-dependent.

Broadly, cells exhibit the ability to recognize and respond to external stimuli. This process is mediated by a coordinated set of extracellular and intracellular interactions that transduce resulting signals into cellular responses^11^. These responses, as a function of dose, define dose-response curves^12^. The dose-response curve is heavily dependent on the type of cell and its internal state^13,14^. Thus, even cells of the same type can respond to the same exposure in a heterogeneous manner^15^. scSeq provides a comprehensive measure of the transcriptome of a cell and captures the inherent variation among cells of the same type. This makes scSeq a useful tool in the study of chemical perturbation of biological systems.

However, a comprehensive cell atlas of chemical perturbations is impossible to assemble, given the vast number of combinations of dose, exposure duration, and cell types^16^. Recently developed resources like the sci-Plex dataset^17^ and the MIX-seq protocol^18^ cover a meaningful but relatively small portion of this space. Algorithms that generalize chemical perturbations across cell state and dose can provide better estimates of the cartography of the chemical perturbation space. In this work we use deep generative modeling to computationally predict cellular response across dose and cell types. We use a class of deep neural networks for dimensionality reduction called autoencoders. Specifically, we use a variational autoencoder^19^ (VAE), which relies on Bayesian priors to encode single cell data into a latent distribution. VAEs have been used to model several technical aspects unique to single cell data, including statistical confounders such as library size and batch effects^20^, and zero inflation^21^. In perturbational single cell biology, VAE models such as scGen^22^ have predicted the response of interferon *β* (IFN-*β*)-treated peripheral blood mononuclear cells (PBMCs). However, when considering more complicated in vivo perturbations, existing models do not consider tissue specific effects when predicting the mean expression of differentially expressed genes (DEGs). VAE’s are uninterpretable in terms of the relationship between latent space and expression prediction. Thus, it is difficult to ascertain which specific genes the model uses to predict differential gene expression after treatment. Thus, there is a need for models that better account for the complexity of in vivo experiments and provide more informative interpretations at the level of individual genes.

Here we propose **single cell Variational Inference** of **Dose-Response** (*scVIDR*), which builds on latent space vector arithmetic when using VAEs to study single cell perturbations (Figure 1). *scVIDR* predicts cell type-specific DEG expression and approximates high-dose experiments better than standard VAEs combined with latent space vector arithmetic (*VAEArith*). We also use *scVIDR* to interpret the latent space using linear models to assess the pathways involved in the single cell dose-response. We accomplish this across several datasets including the dose-response of liver cells to 2,3,7,8 tetrachlorodibenzo-*p*-dioxin (TCDD) in vivo^23,24^, PBMCs treated with IFN-*β*^25^, and a multiplexed dataset of 188 different drug combinations applied to three prominent cancer cell lines (sci-Plex^17^).

**Figure 1.**
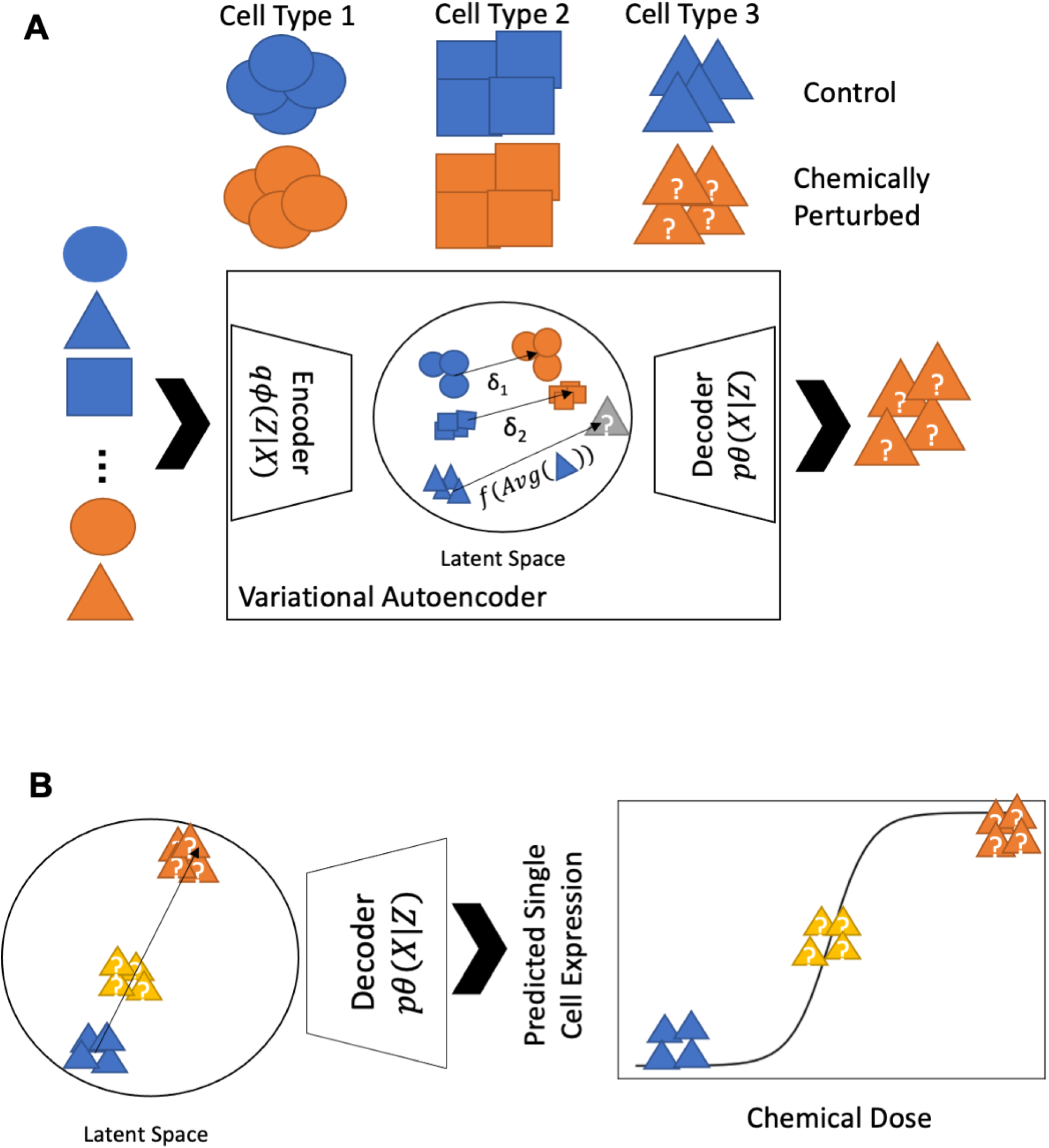
Schematic of scVIDR for prediction of single and multiple doses for some unknown cell type. **A)** Outline of the *scVIDR* model for expression prediction for unknown single dose-response in cell type 3. Training is done using cell types 1 and 2 as input to a variational autoencoder model. The difference between the means of latent representations of the control and treated groups, *δ*_1_ and *δ*_2_, are used as input into a linear regression model. The linear regression model is then used to predict the *δ*_3_ of the unknown cell type. We then use the decoder portion of the model to output the latent space predictions back into gene expression space. **B)** Outline of *scVIDR* for prediction of the unknown response of multiple doses for cell type 3. Log-linear interpolation on *δ*_3_ is used to predict dose dependent changes in gene expression in the latent space. The latent space representations are then projected back into gene expression space using the decoder.

We use data from a single nucleus dose-response experiment in livers from mice gavaged with TCDD as a case study for in vivo dose-response prediction^24,26^. Hepatic responses to TCDD represent an interesting case study as its canonical receptor, the aryl hydrocarbon receptor (AhR), is unevenly expressed along the hepatic lobule, the functional unit of the liver. AhR is more highly expressed in the centrilobular region compared to the portal region (Supplemental Figure 1)^27^. Thus, not only does response to TCDD vary across different cell types in the liver, but also within cell types (such as hepatocytes) along the portal to central axis of the liver lobule^26,28^. To model response variation between cell types, the latent space of the VAE is used to order hepatocytes with respect to their transcriptomic response to TCDD, and thus align all hepatocytes along a “pseudo-dose” axis.

## Results

### scVIDR predicts single-dose, single-cell perturbation expression better than VAEArith

According to the manifold hypothesis, high-dimensional data often lies on a lower dimensional, latent manifold^29^. For single cell data this is a reasonable assumption given that the expression of one gene is often highly dependent on the expression of other genes encoding transcription factors, and is functionally constrained by the process of evolution^30^. Further evidence of this can be seen in the extensive use and success of dimensionality reduction algorithms in the analysis of scSeq data^31^. Lower dimensional representations of single cell data are at the heart of many single cell gene expression analysis methods such as trajectory inference^32^. One method of interest is modeling of the latent manifold using neural networks. These latent manifolds have been shown to simplify complex relationships in single cell gene expression data^33–35^. Specifically, simple vector arithmetic on such spaces can predict in vitro chemical perturbations with high accuracy^16,22^. However, the accuracy of such models when predicting in vivo dose-responses is inconsistent.

We begin by considering a single cell gene expression dataset 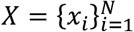 consisting of *N* cells, where *x_i_* represents the expression profile of cell *i*. We assume that gene expression is generated by some continuous random process involving a lower dimensional random variable *z*. The generative process that describes the mapping from *z* to *X* is given by the probability distribution, *p_θ_* (*X*|*z*). Thus, given that we know *X* and not *z*, we would like to approximate the probability distribution that maps *X* to *z*, *p_θ_*(*z*|*X*). Since calculating *p_θ_*(*z*|*X*) is usually intractable we use a neural network, the encoder, to approximate it using a different Gaussian distribution, *q_ϕ_*(*z*|*X*). To map values back from *z* to *X*, we use a second neural network, the decoder, to approximate *p_θ_*(*X*|*z*). In practice, both the encoder and decoder are trained together to minimize the reconstruction error of the decoder and the difference between the prior distribution and the encoder distribution.

We initially developed models for a single-dose chemical perturbation, where we characterize whether a cell has been treated with a set concentration of the chemical of interest with the indicator variable *t* (Figure 1A). We set *t* = 1 for cells that have been treated with the chemical (treatment) and *t* = 0 for cells that have not been treated (control). Our dataset contains *c* cell types within both the *t* = 0 and *t* = 1 groups. Each time a model is evaluated, one treated cell type is withheld from training and used in evaluation. In standard VAE vector arithmetic (VAEArith) the latent space representation of the perturbation of some cell type *A* is approximated by 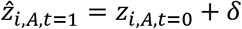. *z*_*i,A,t*=0_ is the latent gene expression representations of cell type *A*^22^, and *δ* is the difference between the means of the treated and control training groups in the latent space. The magnitude and direction of the difference of means between treated and control groups of individual cell types can vary significantly from *δ* (Supplementary Figure 2). Hence, we calculate 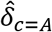, a function of the mean latent representation of the control group of cell type *A*. We approximate this function by training a linear regression model with the other cell types on the latent space (see Online Methods).

We applied this model to the case of a single dose of TCDD administered to mice. Gene expression was measured with single nuclei RNA-seq (snRNA-seq) originating from the mouse liver. We set *t* = 0 for unperturbed gene expression, and *t* = 1 for gene expression perturbed by 30 *μ*g/kg TCDD. The data set covered 6 different liver cell types: cholangiocytes, endothelial cells, stellate cells, central hepatocytes, portal hepatocytes, and portal fibroblasts (Figure 2). Our training set (Figure 2A) consisted of all control and TCDD-treated cell types except for TCDD-treated portal hepatocytes which were used for model evaluation. We evaluated the performance of VAEArith and scVIDR (our method) on the top 5000 highly variable genes (HVGs), and the top 100 differentially expressed genes (DEGs). When predicting the gene expression of portal hepatocytes, each method generated a set of virtual portal hepatocytes. We then computed the average expression of each gene across all cells and compared the average gene expression in predicted cells versus cells derived from snRNA-seq experiments. Across HVGs, the models yielded an *R*^2^ of 0.77 and 0.93, for VAEArith and scVIDR respectively (Figure 2B, C). Across DEGs, the models produced an *R*^2^ of 0.62 and 0.82, for VAEArith and scVIDR respectively (Figure 2C). Examining the expression of an individual gene known to robustly respond to TCDD perturbation, *Cyp1a2*, scVIDR predicted its mean expression better than VAEArith (Figure 2D)^28^. Continuing the evaluation across all cell types (Figure 2F), leaving out one cell type perturbation at a time as described above for portal hepatocytes, our model outperformed VAEArith (with p-value < 0.001, one sided Mann-Whitney U Test) both when evaluated on HVGs and DEGs.

**Figure 2.**
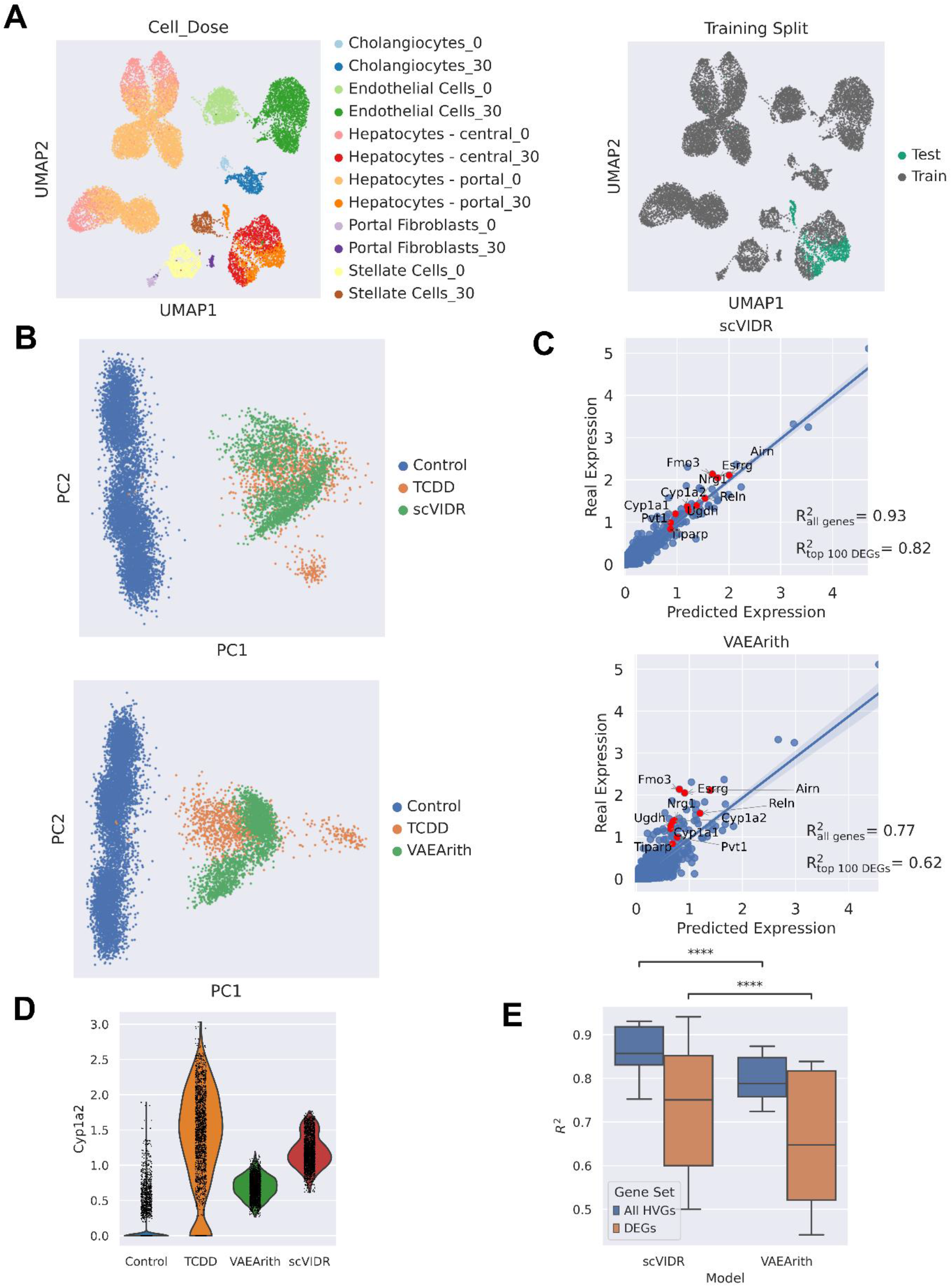
Prediction of in vivo single cell gene expression of portal hepatocytes from mice treated with 30 *μ*g/kg of TCDD. **A)** UMAP projection of the latent space representation of control and treated singlecell gene expression. In **A)** each cell type is represented by a different color, the dose administered and by the train-test split for model training. In the example in the figure, TCDD treated portal hepatocytes were used as a test set. **B)** PCA plots of predicted portal hepatocytes responses following treatment with 30 μg/kg of TCDD using VAEArith, and *scVIDR*. **C)** VAEArith and scVIDR prediction versus real expression regression plot. Each point represents the mean expression of a particular gene. Red points represent the top ten differentially expressed genes. **D)** A violin plot of real and predicted Cyp1a2 expression by VAEArith and scVIDR. **F)** Boxplot of *R*^2^ of predicting across all liver cell types treated with 30 μg/kg of TCDD. Calculation of the mean *R*^2^ across all highly variable genes (blue). Calculation of the mean *R*^2^ across the top 100 differentially expressed highly variable genes (orange). Prediction performance distributions were compared using one sided Mann-Whitney U test which showed p-values of less than 1e-4 for higher overall performance in scVIDR.

We had similar results for Interferon *β* (IFN-*β*)-treated peripheral blood mononuclear cells -PBMCs (Supplementary Figure 3)^25^. Here *t* = 1 for PBMCs treated with IFN-*β*, and *t* = 0 for untreated PBMCs (Supplementary Figure 3A). Across HVGs, the models yielded *R*^2^ values of 0.92 and 0.97, and across DEGs, *R*^2^’s of 0.87 and 0.92, for VAEArith and scVIDR, respectively (Supplementary Figure 3C). When accuracy was assessed for all cell types, scVIDR significantly outperformed VAEArith (Supplementary Figure 3D).

### scVIDR accurately predicts the transcriptomic response for multiple doses across cell types

Next, we predicted the response for multiple doses of TCDD (Figure 1B). Here, *p* is equal to the magnitude of the perturbation, which in our case is equivalent to the dose. Thus, *p* = 0 represents expression at dose 0 and *p* = 30 represents expression at dose 30, where the dose is in units of *μ*g/kg in Figure 3 and nM in Supplementary Figure 4. As with the single-dose case, we train the model on the dose-response data for all cell types except one, for which only the *p* = 0 condition is kept. We calculate the 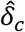 (Online Methods, Figure 3A) which is the estimated difference of means between the highest dose and the untreated groups. Other doses are then calculated on the latent space by interpolating log-linearly on the 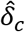. Finally, those latent space representations are decoded back into gene expression space using the decoder portion of scVIDR.

**Figure 3.**
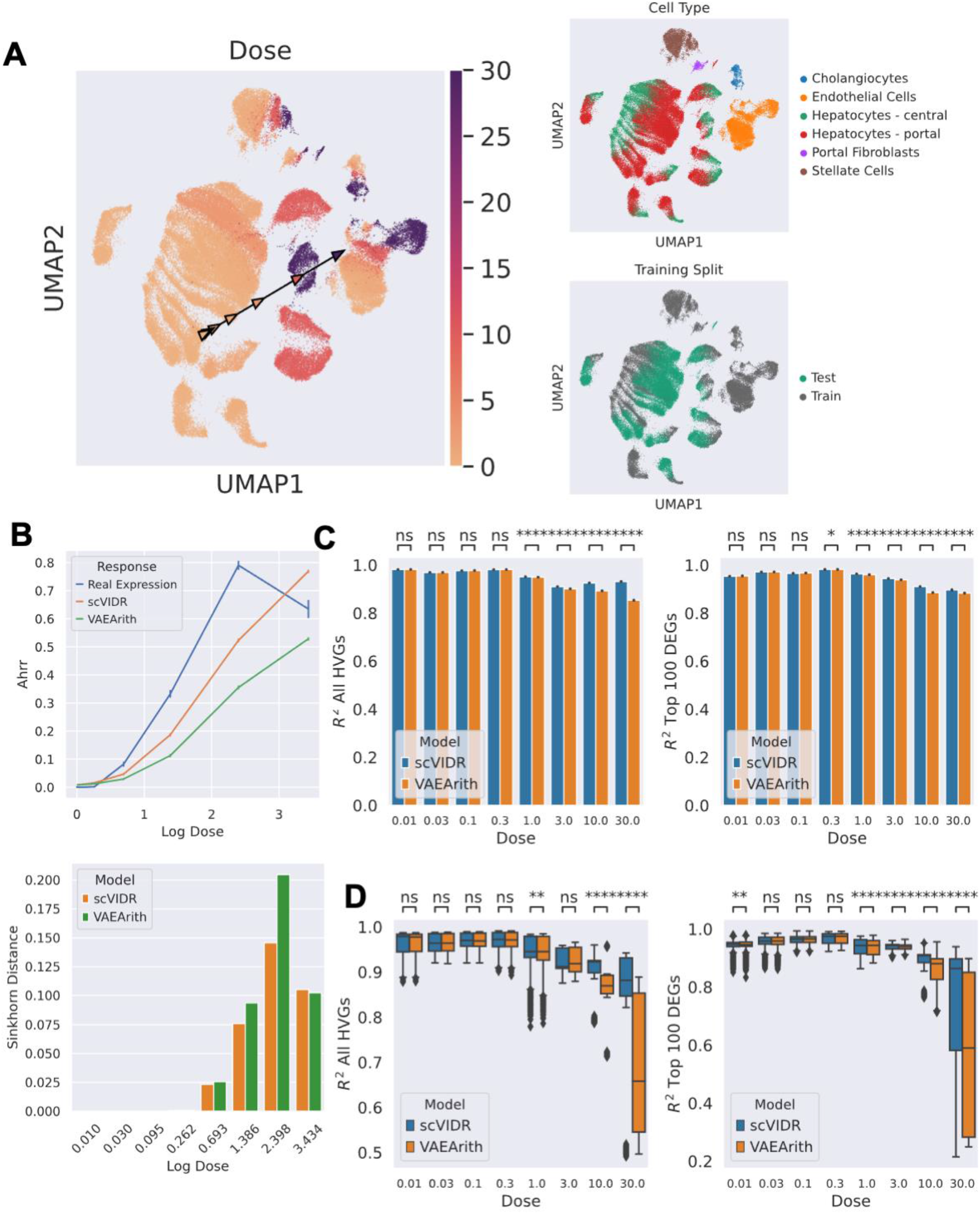
Prediction of in vivo singe cell expression of the TCDD dose-response in portal hepatocytes from mouse liver tissue. **A)** UMAP projection for the latent space representation of single cell expression across TCDD dose-response. Cells are colored by dose (*μ*g/kg), cell type, and training split. Arrows on UMAP represent a *δ* calculated on UMAP space, with each arrowhead representing a specific dose denoted by its color. **B)** Dose-response prediction for *Ahrr* using scVIDR, and VAEArith. The differences between the predicted and true distribution of Ahrr at each dose are measured via the Sinkhorn distance. **C)** Bar plots of the *R*^2^ of the gene expression means in portal hepatocytes for all highly variable genes and the top 100 differentially expressed genes. **D)** Box plot of the distribution of *R*^2^ scores across all cell types in liver tissue.

We analyzed a mouse liver snRNA-seq dataset that included 8 doses (p = [0.01, 0.03, 0.1, 0.3, 1.0, 3.0, 10, 30]) of TCDD and a control (p = 0) in *μ*g/kg (Figure 3). scVIDR outperforms VAEArith in approximating expression across the dose-response of TCDD in mouse liver. We used the mean *R*^2^ score across all evaluated genes as our performance metric (Figure 3B). scVIDR significantly out-performed VAEArith at predicting HVGs and DEGs for doses > 0.3 *μ*g/kg (Mann-Whitney One-Sided U test p < 0.001). scVIDR predicts the important TCDD receptor repressor gene, *Ahrr*, at doses 1, 3, and 10 *μ*g/kg in portal hepatocytes better than VAEArith (Figure 3C). When looking across all possible training scenarios (leaving one cell type out) scVIDR significantly outperformed VAEArith only at the highest doses of 10 and 30 *μ*g/kg on prediction of all HVGs (Figure 3D). When predicting on just the DEGs, scVIDR significantly outperformed VAEArith for doses > 0.3 *μ*g/kg (Figure 3E).

We used scVIDR to predict the effects of a test set of 37 drugs out of 188 treatments in the sci-Plex dose-response data^17^ at 24 hours for A549 cells (Supplemental Figure 4A). scVIDR was trained on all data (all drugs and doses) in K562 and MCF7 cells. The model was also trained on the remaining 151 drugs in A549 cells not used in validation, as well as the vehicle data (Supplemental Figure 4A). The dose-response for the 37 drugs was predicted as above by first calculating the 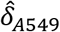 between the control and highest dose for a particular drug and log linearly interpolating along the 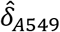 in order to predict the intermediate doses. We evaluated predictions made by scVIDR at the gene, drug, and drug pathway level. For the drug Belinostat, a histone deacetylase inhibitor, scVIDR improves on predictions of differentially expressed genes such as *MALAT1* relative to VAEArith (Supplemental Figure 4B). When predicting gene expression of the DEGs in Belinostat treated A549 cells, scVIDR also significantly outperformed VAEArith on all doses (Supplemental Figure 4C). On predicting the DEGs of all drugs with the same mode of action as Belinostat (Epigenetics), scVIDR similarly outperformed VAEArith on all doses (Supplemental Figure 4D). Finally, when looking across all 37 drugs in the test dataset, we were able to predict the expression of DEGs significantly better than VAEArith on average for the 3 highest doses of 100, 1,000, 10,000 nM (Supplemental Figure 4E).

### Regression on the latent space infers the relationship between predicted gene expression and 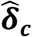

Insight into model decisions can provide information regarding proper model usage and pitfalls. It would be useful to identify which genes and pathways are associated with scVIDR’s prediction; however, standard VAEs are not easily interpretable. To interpret scVIDR’s predictions, we approximate the function of the decoder with linear regression (see Online Methods).We take inspiration from the use of principal component analysis (PCA) in scSeq^36^ and the development of the linear decoded variational autoencoders (LDVAE)^37^. PCA is a linear transformation that projects the data onto a lower dimensional (latent) space while retaining as much variance as possible. This transformation is represented by a linear weight matrix, *W_pca_*, with dimensions *m x g* where *m* is the number of latent variables, and *g* is the number of genes. We can understand each principal component as a linear combination of genes. This allows us to assess the relationship between genes and a direction in latent space.

In a VAE, the mapping from the latent space to the gene space is done by the decoder which, unlike the inverse of PCA, is non-linear. In LDVAEs, however, the decoder portion of the VAE is a linear regression layer and thus the weight matrix of this layer, *W_ldvae_*, describes a linear relationship between direction in the latent space and gene prediction^37^.

However, interpretability comes at the expense of model accuracy. LDVAEs have higher reconstruction error than standard VAEs on single cell data^37^. Similarly, using PCA and vector arithmetic to predict scSeq perturbations performed poorly compared to VAEArith^22^. As a result, one would like to try and interpret the latent space of a standard VAEs. We present an approach to interpret the VAE’s latent space using sparse regression.

We take an alternative approach to LDVAEs in which we instead approximate the non-linear function of the decoder in a standard VAE using sparse linear regression (Figure 4A). Sparse regression methods like LIME have been used to interpret complex models^38^. We specifically use sparse linear ridge regression, given that teach gene has a non-zero contribution to each latent variable and gene weights are distributed parsimoniously. This gives us a linear transformation matrix, 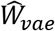, that approximates the function of the decoder.

**Figure 4.**
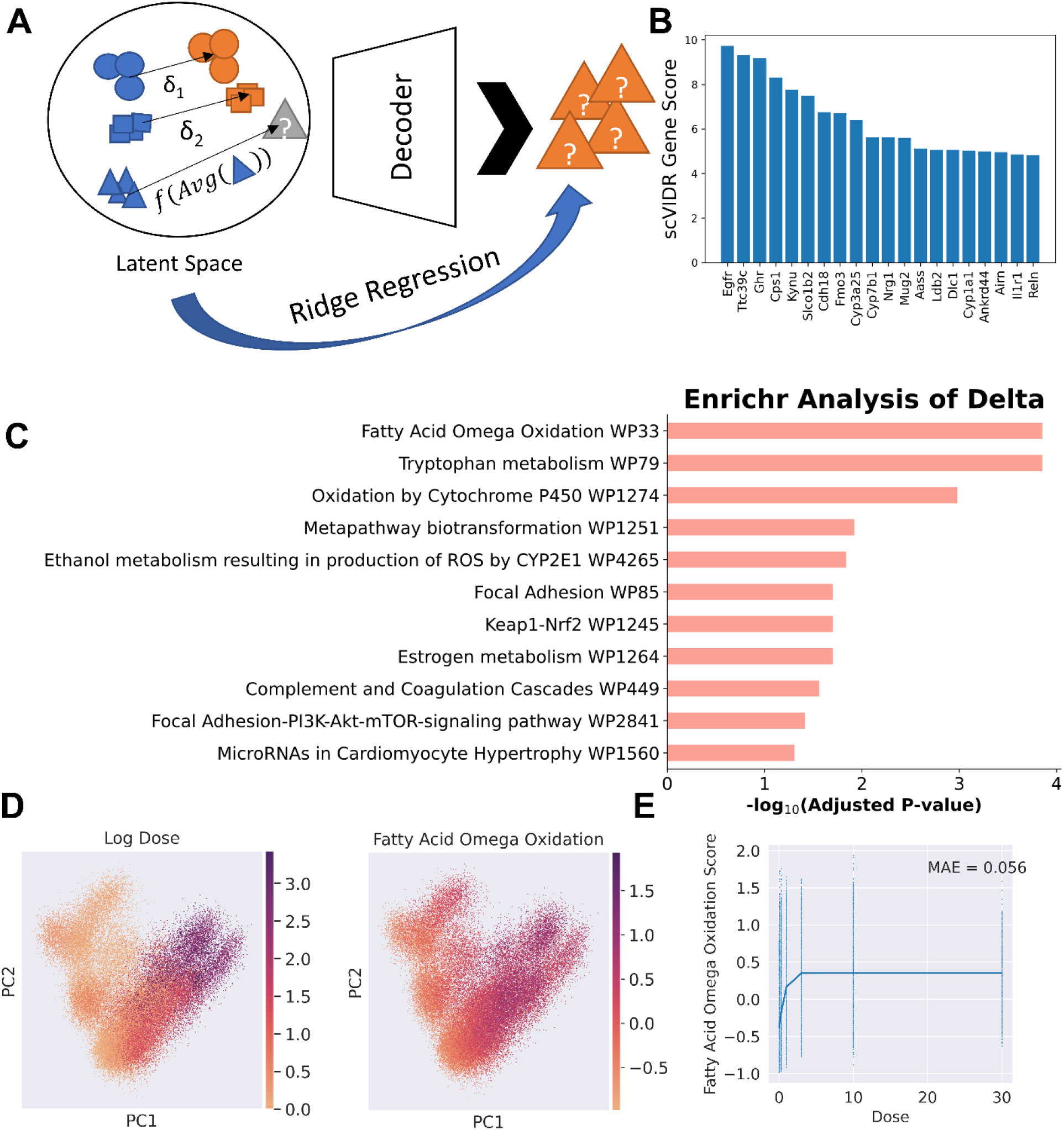
Interrogation of VAE using ridge regression in portal hepatocyte prediction. **A)** Schematic of calculating latent dimension weights using ridge regression. **B)** Bar plot of top 20 genes with the highest scVIDR genes scores. **C)** Enrichr analysis of the top 100 genes with respect to the scVIDR gene scores. Bar plot of adjusted p-values from statistically significant (adjusted p-value < 0.05) enriched pathways from the WikiPathways 2019 Mouse Database. **D)** PCA projection of single cell expression data colored by log dose and fatty acid oxidation pathway score. **E)** Logistic fit of median pathway score for each dose value. MAE - mean absolute error.

We use this weight matrix to interrogate the relationship between predicted gene expression and 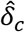. The span of 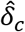 is simply a direction in scVIDR’s latent space. The importance of 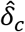 to each gene’s predicted expression is the sum of the latent dimensional components of 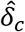 multiplied by the gene’s corresponding latent dimensional weight from 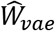. In matrix form:

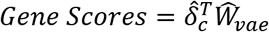

In practice, we found that normalizing the weight matrix by its L2 norm gives better insights when interpreting the model (see Online Methods).

We utilize a trained VAE model where portal hepatocytes were left out of training and the 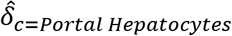 was approximated (Figure 4 B-D). Gene scores for 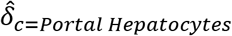 were calculated as described above. The genes with the top 20 highest genes scores included well established markers of TCDD-induced hepatotoxicity such as genes from the cytochrome P450 family (Figure 4B)^28^. To see whether this relationship extended to pathways involved in TCDD-induced hepatotoxicity, we performed Enrichr analysis^39^ using the 2019 WikiPathways database^39^ on genes with the top 100 gene scores (Figure 4C). Among the top enriched terms, we found the hallmarks of hepatic response to TCDD in mice, such as oxidation by cytochrome P450^40^, fatty acid omega oxidation^41^, and tryptophan metabolism^42^. To derive the relationship between the actual doses and the gene pathways, the genes with the top 100 gene scores that were in “Fatty Acid Oxidation” from WikiPathways was used in calculating enrichment scores for each cell using Scanpy^43^. A sigmoid function was fit to the median enrichment score in each dose (see Online Methods). We observed a small mean absolute error in our model and thus concluded that there was a sigmoidal dose-response relationship for the gene set generated by Enrichr (Figure 4 D, E).

### Pseudo-dose captures zonation in TCDD hepatocyte response

In single cell analysis of developmental trajectories, it is useful to order cells with respect to a latent time course, termed “pseudo-time”. This is because cells develop at different rates due to natural variations among themselves and their environment. This ordering is usually done using algorithms such as Slingshot^44^ and Monocle^45^. In pharmacology and toxicology, we experience a similar problem as cells of the same type have variable sensitivities to the same toxicant. Hence we propose to order cells in terms of a latent dose. We call this ordering of cells a “pseudo-dose”.

Working off the assumption that *δ_c_* (see Online Methods) is the axis of perturbation in latent space, we orthogonally project the latent representation of each cell to the *span*(*δ_c_*) to obtain a scalar coefficient for each cell along *δ_c_* (Figure 5A, B). We use this scalar coefficient as the pseudo-dose value for each cell.

**Figure 5.**
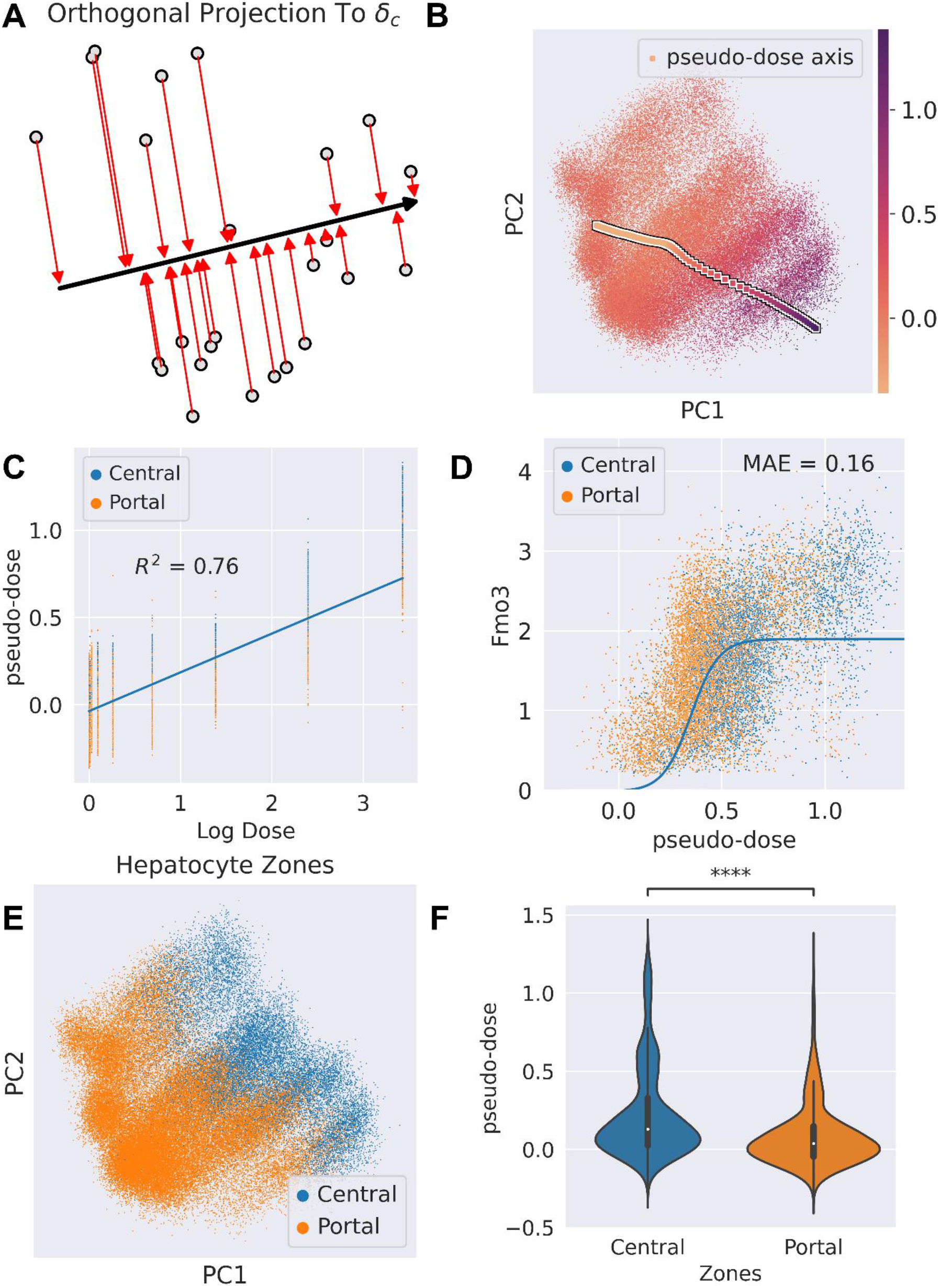
Pseudo-dose ordering of hepatocytes across TCDD dose-response. **A)** Schematic diagram of assigning pseudo-dose values to hepatocytes by orthogonally projecting each cell in latent space to the span of the *δ_c_*. **B)** PCA projection of hepatocytes colored by assigned pseudo-dose values. The arrow markers represent the pseudo-dose axis calculated by the *δ_c_*. **C)** Regression plot of pseudo-dose vs. log transformed dose **D)** Plot of pseudo-dose vs. *Fmo3* expression. Associated logistic fit (solid blue line) and associated mean absolute error annotated as “MAE”. **E)** PCA projection of hepatocyte expression colored by assigned hepatocyte zone in liver lobule. **G)** Violin plot of the distribution of pseudo-dose values in the central and portal zones of the liver lobule. Central hepatocytes exhibit a higher pseudo-dose on average than portal hepatocytes (Mann-Whitney Single Sided U-Test p-value < 1e-4).

To test whether these pseudo-dose values capture the latent response across cell types, we distinguished between the portal and central regions of the liver lobule. Zonation of the lobule not only defines differences in hepatocyte gene expression along the portal to central axis, but also their metabolic characteristics ^46^. Thus, we expect that the two zones will exhibit different sensitivities to TCDD. The pseudo-dose correlated well with the actual dose administered to the hepatocytes with an *R*^2^ = 0.76 (Figure 5C). We also found that pseudo-dose displayed a sigmoidal relationship (see Online Methods) between the expression of differentially expressed genes such as Fmo3 (Figure 5D). Finally, we found the pseudo-dose to be statistically higher on average in the central hepatocytes versus portal hepatocytes (Figure 5E). This is consistent with liver biology as central hepatocytes respond more strongly to treatment due to TCDD sequestration^47^, and higher AhR expression levels in the centrilobular zone^27^.

## Discussion

Mapping the combinatorial space of single-cell perturbation is important to toxicology and pharmacology to facilitate the generalization of drug or toxicant effects across several domains. Computational modeling allows researchers to use current large-scale databases to predict new perturbations to scSeq data. We have demonstrated an improvement to such modeling using VAEs. These improvements include highly correlated prediction of cell type specific effects in mouse liver, PBMCs and A549 cells. We also modeled a latent response for mouse hepatocytes using pseudo-dose and interrogated the VAE to predict dose-dependent perturbations in portal hepatocyte pathways. We show that deep generative modeling can be used to model complex perturbations in single-cell gene expression data from several different datasets.

When evaluating the model in the mouse liver, scVIDR performed better on the cell types most sensitive to TCDD, e.g., hepatocytes and endothelial cells (Supplementary Figure 5A, C, D). When looking at cell types less sensitive to TCDD, the model often underestimated the expression of differentially expressed genes (Supplementary Figure 5E). This is likely a result of a combination of factors including the similarity of the treatment to the control data (Supplementary Figure 5A), smaller control cell populations (Supplementary Figure 5B), and the overall low expression of highly variable genes (Supplementary 5E). Thus, we believe the VAE has less information to predict differential gene expression for these cell types. Our model improves on this problem with respect to VAEArith for most cell types in the liver (with the exception of stellate cells and cholangiocytes at higher doses). Results from sci-Plex imply that incorporating scSeq data from livers treated with other compounds could improve these predictions, as the model would have more information on different liver responses.

In the sci-Plex dataset, prediction of certain drugs with epigenetic mode of actions produced the poorest prediction scores (Supplementary Figure 6). This is because scSeq data provides no information regarding epigenetic modifications (e.g., chromatin accessibility, histone marks, and DNA binding proteins). Integration with epigenetic scSeq data such as single cell ATAC-seq could help to predict such responses with higher accuracy.

When looking to the future of generative modeling in chemical-induced perturbation of gene expression, a problem domain of interest is time-dependent drug effects. Chemical exposures are not only a function of concentration, but also of time^48^. Dose-Time-Response analysis is central to risk assessment in clinical settings^49^. Predicting the response not only as a function of amount of drug, but also as a function of the time the drug is within a patient’s system and the time of day at which the drug was administered would allow for more effective and safer dosing regimens^49,50^. Developmental state can also be impacted by chemical perturbation. An example of this is the inhibition of B-cell lymphopoiesis by TCDD^51^. The latent space could be useful for analyzing a simplified model of the dynamics of developmental systems and how they change with chemical perturbation. Principal component analysis for dimensionality reduction has been used in this area for successful cellular fate prediction during hematopoiesis^52^.

Taken together, our tool facilitates dose-response predictions for a particular drug in a specific cell type using the response of other cell types. Future directions outlined above can yield powerful tools in the realm of pharmacology and toxicology. As more data becomes available on single cell chemical perturbations, generative modeling and other forms of dimensionality reduction can yield insights into the underlying manifold of gene expression and how different classes of chemicals act on that manifold. Discovery of the properties of this manifold will allow for more generalizations to be made about the physiology of tissues and understudied chemical perturbations.

## Online Methods

### Single cell expression datasets and preprocessing

All the datasets except the PBMC IFN-*β* treatment dataset were collected and processed uniformly from raw count expression matrices. Single nuclei RNA-seq of C57BL6 mouse liver administered with different doses of TCDD is obtained from Nault et al. This data can be found under GSE184506 in the Gene Expression Omnibus (GEO)^24^. Mice in this dataset were administered, sub-chronically, a specified dose of TCDD via oral gavage every 4 days for 28 days. In our analysis, all immune cell types were left out, as immune cells are known to migrate from the lymph to the liver during TCDD administration^23^. Thus, there is a small size for the immune cell populations in the low-dose datasets versus the higher doses. The *sci-Plex* dataset is obtained from Srivatsan et al and can be found under GSE139944 from GEO^17^. The cell expression vectors are normalized to the median total expression counts for each cell. The cell counts are then log transformed with a pseudo-count of 1. Finally, we select the top 5000 most highly variable genes to do our analysis on. The preprocessing was carried out using the *scanpy.pp* package using the *normalize_total*, *log1p*, and *highly_variable* functions^43^. PBMC data from Kang et al^25^ was accessed as a processed dataset from Lotfollahi et al^22^. The PBMC dataset can be found under GSE96583 in GEO.

### Variational autoencoder

Variational Autoencoders^19^ aim to estimate the posterior probability function, *p_θ_*(*z*|*X*) of a latent process, *z*, given a set of observations, *X*. By Bayes theorem we can calculate the posterior probability:

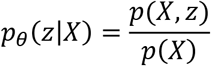

However, calculating the posterior in this way is intractable due to the difficulty in computing the marginal distribution:

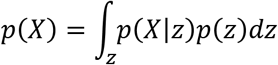

Thus, we instead aim to approximate *p_θ_*(*z*|*X*) by minimizing the Kullback-Leibler (KL) divergence between *p_θ_*(*z*|*X*) and some gaussian distribution, *q_φ_*(*z*|*X*), whose parameters, *ϕ*, are calculated by a neural network. This neural network is termed the encoder.

We can calculate the KL divergence as:

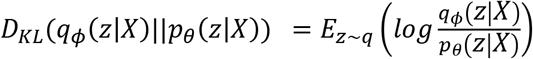

Substituting *p_θ_*(*z*|*X*) with Bayes theorem and rearranging the terms:

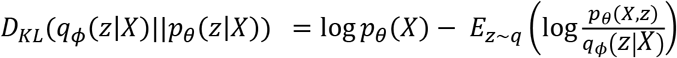

The second term in the equation above is also known as the evidence lower bound, or ELBO. Since the KL divergence must be a positive value, we can minimize it by maximizing the ELBO. We can rewrite the ELBO as:

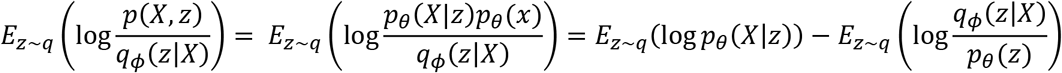

Since the second term in the equation above is equivalent to the definition of the KL Divergence we can rewrite the ELBO as:

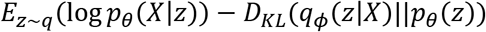

The first term is known as the reconstruction error. This maximizes the likelihood that we will generate values from the latent space that match our observations. The second term is the KL divergence term, which is predicated by the KL divergence between the distribution estimated by the encoder and the prior distribution which is a standard normal multivariate distribution. We can now construct our objective function, *L*(*θ*, *ϕ*), as the following:

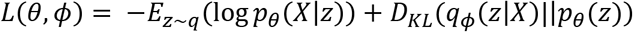

Unfortunately, naively trying to take the gradient with respect to *ϕ* to minimize this function will result in highly variable gradients and thus variable training results. To fix this we instead utilize the reparameterization trick, where we sample from the latent space such that *z* is deterministic with respect to some noise variable, *ϵ* ~ *N*(0, *I*), or:

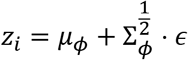

Where, *μ* is the mean vector and Σ is the covariance matrix of the inferred distribution.

### Implementation of variational autoencoder

All code in this manuscript is implemented in the Python programming language. The scVIDR model is built on the python package, scGen v. 2.0.0^22^. Here we modify the model to accommodate predictions of the dose-response, linear regression on the latent space, pseudo-dose calculations, and approximations of the gene importance in chemical perturbations.

#### Calculation of the *δ_c_* for single and multiple dose predictions

The *δ*, as defined by Lotfollahi et al^22^, is the difference between the mean latent representations of the treated (t=1) and untreated (t=0) conditions.

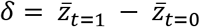

Where, 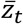 is the mean latent representation for treatment *t* in the dataset.

We can calculate a cell type specific *δ*_*c*=*A*_ for some cell type, *A*, by taking the difference between the mean latent representations of the treated and control groups, or:

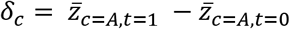

If we want to estimate a *δ_c_* for some type of cell type *B* based on 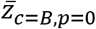 and where 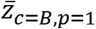 is unknown, we can approximate a function based on 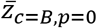, or:

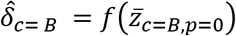

Where we approximate the above function using all other existing cell types in the dataset as input to ordinary least squares regression as implemented by *LinearRegression* function in the *sklearn.linear_model* package^53^.

To predict the latent representation for a response at some dose, *d*, we interpolate log-linearly on 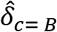 such that for each latent cell in our prediction, *Z*_*i,c,p*=*d*_:

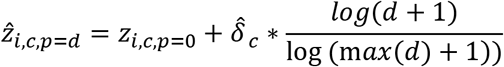

### Evaluating model performance

Performance of the model on the prediction task is the same as that in Lotfollahi et al^22^. We quantified performance using the *R*^2^ value for mean gene expression for each gene across all cells. The *R*^2^ was calculated using the *linregress* function from the *scipy.stats* package^54^. We compared the DEGs are selected using the *rank_gene_groups* from the *Scanpy* package and taking the top 100. Models were compared on the same prediction in which we resample 80% of the cells in the cell type we are predicting 100 times. Resampling is done using the *choice* function from the *numpy.random* package^55^.

Statistical significance was determined by the one-sided Mann-Whitney U test as it is implemented by *mannwhitneyu* function from the *scipy.stats* package. We considered p-values less than 0.001 as statistically significant.

Distances were used to establish relationships between distributions and vectors. Cosine distance was calculated using the *cosine* function in the *scipy.spatial.distance* package. The Sinkhorn distance was calculated using the *SampleLoss* class in the *geomloss* package^56^.

### Inferring feature level contributions to perturbation prediction

In PCA, we perform an orthogonal linear transformation on the data, such that our projected data preserves as much variance as possible. It is known that the solution to this maximization problem is to project the data onto the eigenvectors of the covariance matrix or:

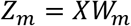

Where *X* is the mean-centered scRNA-seq expression matrix, *W_m_* is the eigenvectors corresponding to the *m* highest eigenvalues of the covariance matrix of *X*, and *Z_m_* represents the *m*-dimensional projection of the data onto its principal components. We can see from this formula that *Z_m_* is calculated as a linear combination of weights and gene expression, and thus there is a linear relationship between genes and the principal components. We can exploit this fact and calculate a loading for each gene with each corresponding eigenvector by taking the product of the eigenvector and its corresponding eigenvalue or:

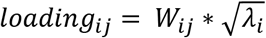

Where *W_ij_* is the *j^th^* value (corresponding to gene j) of the *i^th^* eigenvector and *λ_i_* is the eigenvalue for the *i^th^* eigenvector. These loadings represent a normalized score of the relationship between a gene’s expression and a particular principal component. These loadings are also directly proportional to the actual correlation between the gene’s expression and the principal component of interest.

It can be shown that PCA and autoencoders with a single hidden layer (with size less than the observations) and a strictly linear map are nearly equivalent^57^. We can project principal components back into expression space using the following function:

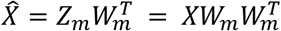

Additionally, we note that PCA is a solution to the minimization of the reconstruction error:

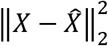

We find similarly that the loss function that we try to optimize in the autoencoder we described above is:

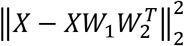

Where *W*_1_ is the weights of the hidden layer, and *W*_2_ is the weights of the final layer of the autoencoder. In effect, we can see that the autoencoder described above can approximate the loadings of a principal component analysis using *W*_2_.

The reconstruction error for a standard VAE with the assumption that the observations are a multivariate Gaussian is:

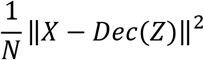

Where *N* is the number of samples, *Dec*(*Z*) is the function of the decoder neural network, and *Z* is the transformation by the encoder of the observations onto the latent space. In an LDVAE, the *Dec*(*Z*) is replaced with single layer with linear transfer operators such that the reconstruction error is the following:

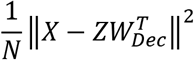

In which *W_Dec_* is the linear weights of the decoder. These weights give us an approximation of the contributions of individual genes to the dimensions of the latent space. We can interpret *W_Dec_* as a loadings matrix by which we can interpret the latent dimensions of the LDVAE.

To approximate feature contributions to predicting the perturbation in scVIDR, we train a ridge regression model. We then take the decoder portion of our model and sample 100,000 points from the latent space and generate their corresponding expression vectors. This will be our training dataset for a ridge regression. We then train the ridge regression using the *Ridge* class from the *sklearn.linear_model* package. We can describe the loss of our ridge regression as:

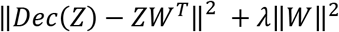

Where *Z* are the sampled points from the latent space, *ZW^T^* are the approximation of the predicted gene expression vectors, and *W*, is a m x n matrix where m Is the number of genes and n is the number of latent dimensions. We divide *W* using the ║*W*║_2_ to normalize for the effect of over expressed genes. We then calculate the gene scores by taking the dot product of normalized *W* and *δ_c_*, or:

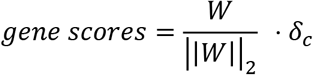

We use these gene scores to order genes for *Enrichr* pathway analysis with the *gseapy* package. Scores for each pathway were calculated using *score_genes* function from *scanpy.tl* package with the genes sets derived from the *Enrichr* results.

#### Calculating the pseudo-dose values

We can order each cell, *x_i_*, with respect to the variable response of *X_i_* to the chemical by taking the latent representation, *z_i_*, and orthogonally projecting it onto *L* = *span*(*δ*):

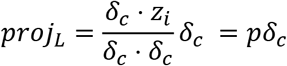

The scalar multiple of *δ*, *p*, is the pseudo-dose value for *x_i_*.

#### Regression of sigmoid function for evaluating dose-response relationships

To establish whether a standard dose-response relationship existed between the top pathways inferred by *Enrichr* and between the pseudo-dose and gene expression, a logistic function of the form:

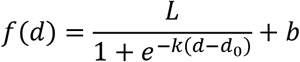

Where d is the dose or pseudo-dose. The parameters of the function above were fit to the output variables (median enrichment score and Fmo3 normalized expression) using Levenberg-Marquardt algorithm implementation in the *curve_fit* function in the *scipy.optimize* package. The regression was evaluated using the mean absolute error metric implementation in the *mean_absolute_error* function in the *sklearn.metrics* package.

## Data Availability

All data used in the manuscript is publicly available and referenced in the manuscript.

## Code Availability

The code for the software and for reproducing the figures is available at TBA.

## Acknowledgments

This work was supported by the National Human Genome Research Institute R21 HG010789 to T.Z. and S. B. O. K. is supported by the National Institute of Environmental Health Sciences of the National Institutes of Health under awards number T32 ES007255. T. Z. and S.B. are partially supported by the USDA National Institute of Food and Agriculture, Michigan AgBioResearch. The content is solely the responsibility of the authors and does not necessarily represent the official views of the National Institutes of Health. This work was supported in part through computational resources and services provided by the Institute for Cyber-Enabled Research at Michigan State University.

## Author contributions

S.B. conceived and conceptualized the research with contributions from O.K.

R.N. performed the *in vivo* experiments and generated the TCDD dose-response experimental dataset. O.K. downloaded and preprocessed the sci-Plex and PBMC datasets, implemented and evaluated models, and generated figures. O.K. and D.F. developed model interpretation metrics. R.N. and T.Z. contributed to the biological interpretation model. D.M. and D.F. helped with the bioinformatics analyses. R.N., T.Z., D.M., and D.F. all helped with writing results. All authors contributed and reviewed the manuscript.

---

**Supplementary Figure 1.**
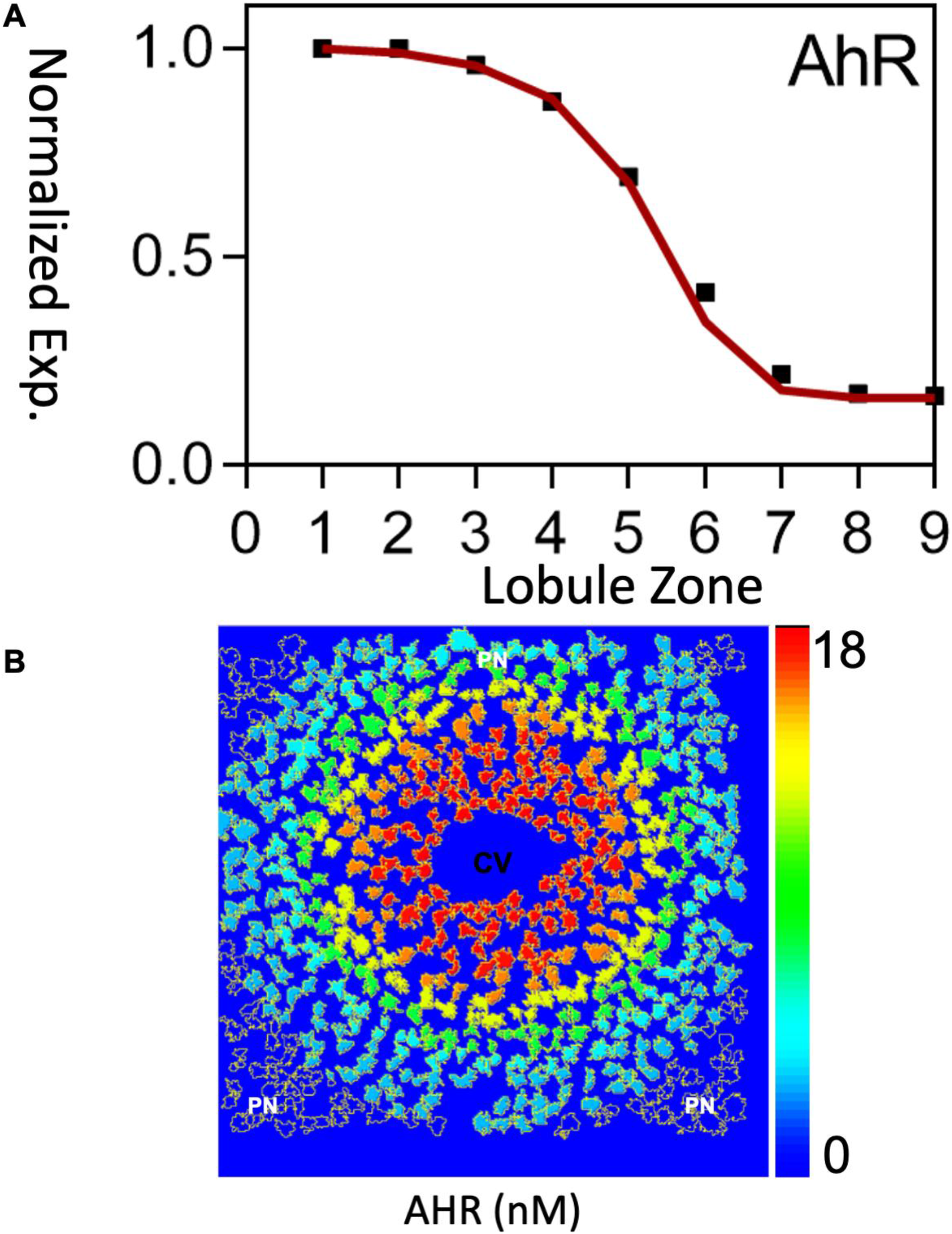
Expression of AhR across liver lobule. **A)** A zonal expression profile of normalized expression as described by Yang et al.^28^ and Halpern et al.^58^ Zone 0 represents the level of AhR expression in hepatocytes closest to the central vein. Zone 9 represents the level of AhR expression closest to the portal vein. **B)** A single cell resolution image of the liver lobule generated by Halpern et al. with expression levels represented by color from Yang et al. The central vein is denoted by “CV” (black) with the portal triad denoted by “PN” (white).

**Supplementary Figure 2.**
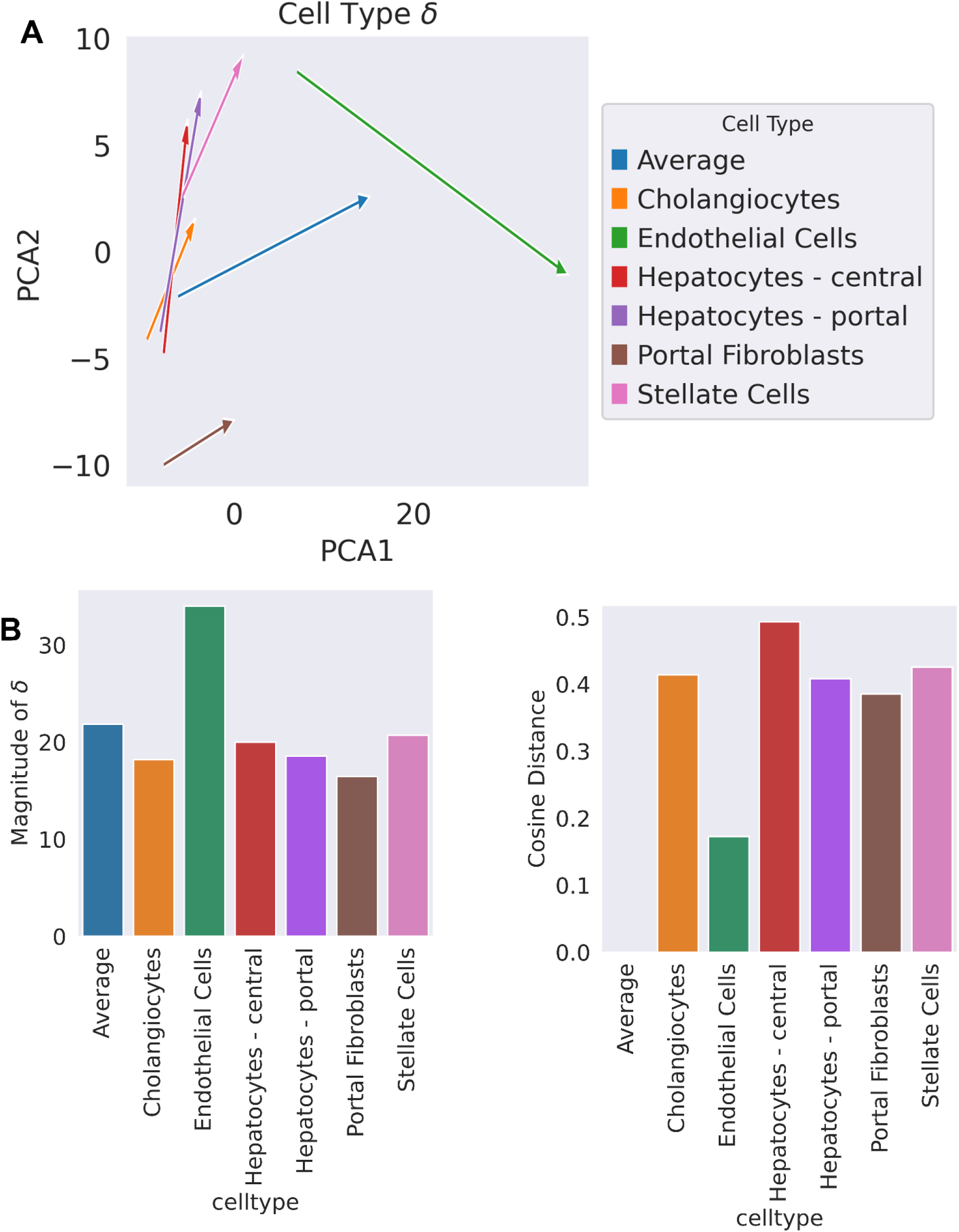
*δ_c_ s* deviate from *δ* with respect to their direction across most cell types in the TCDD mouse liver snRNA-seq dataset. **A)** A PCA visualization of the calculated *δ_c_*s for a VAE trained without portal hepatocytes. “Average” denotes the standard *δ* as defined by scGen^22^. **B)** Bar plots of the magnitude of the *δ* and other individual *δ_c_’s*, and *δ_c_* cosine distance from the *δ*. A cosine distance of 0 represents a *δ_c_* in the same direction as *δ*, of 1 represents a *δ_c_* orthogonal to *δ* and of 2 represent a *δ_c_* in the opposite direction as *δ*.

**Supplementary Figure 3.**
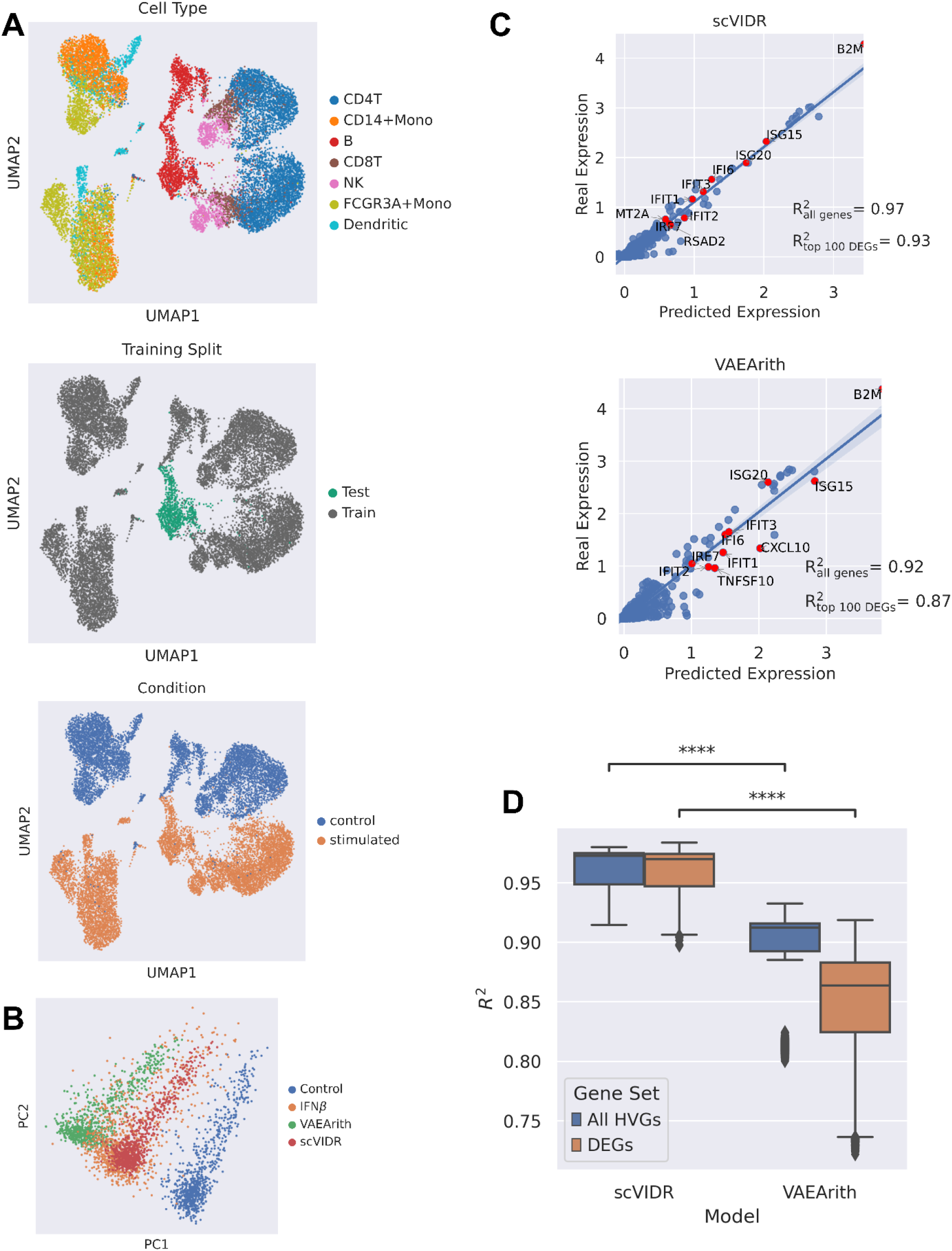
Prediction of in vitro response of B-cells to IFN*β*. **A)** UMAP of latent space of treated and untreated single-cell expression. UMAP plots are colored by cell type, training split, and condition, respectively. **B)** PCA plot of VAEArith and scVIDR predictions of B-cell expression after IFN*β* treatment. **C)** VAEArith and scVIDR prediction versus experimental expression data regression plot. Each point represents the mean expression for a particular gene. Red points represent the top ten differentially expressed genes. **D)** Boxplot of *R*^2^ scores across all tissues in the PBMC treated dataset. Prediction of all highly variable genes (blue), and top 100 differentially expressed genes (orange).

**Supplementary Figure 4.**
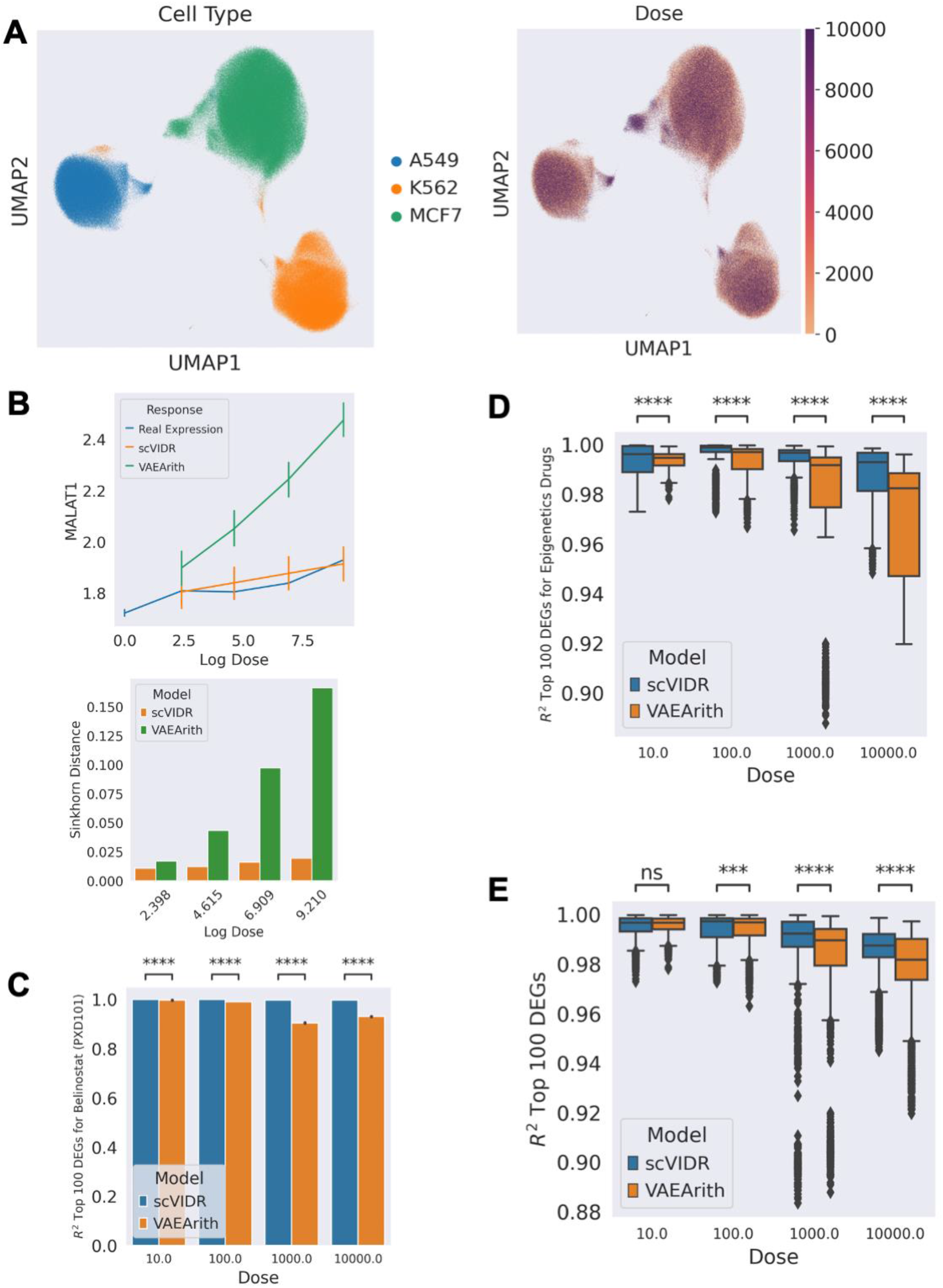
Prediction of in vitro dose-response of A549 cells to different drug treatments. **A)** UMAP of the latent space of single-cell expression colored by cell type and dose (nM) respectively. **B)** Prediction of the dose-response of MALAT1 in response to Belinostat treatment of A549 cells. The differences between the predicted and true distribution and of MALAT1 at each dose are measured via the Sinkhorn distance. **C)** Bar plot of prediction performance of the dose-response of Belinostat administered to A549 cells on the top 100 differentially expressed genes **D)** Boxplot of prediction performance of the top 100 differentially expressed genes for the A549 dose-response in all test dataset epigenetic pathway drugs. **E)** Boxplot of prediction performance of the top 100 differentially expressed for the A549 dose-response in all 37 test dataset drugs.

**Supplementary Figure 5.**
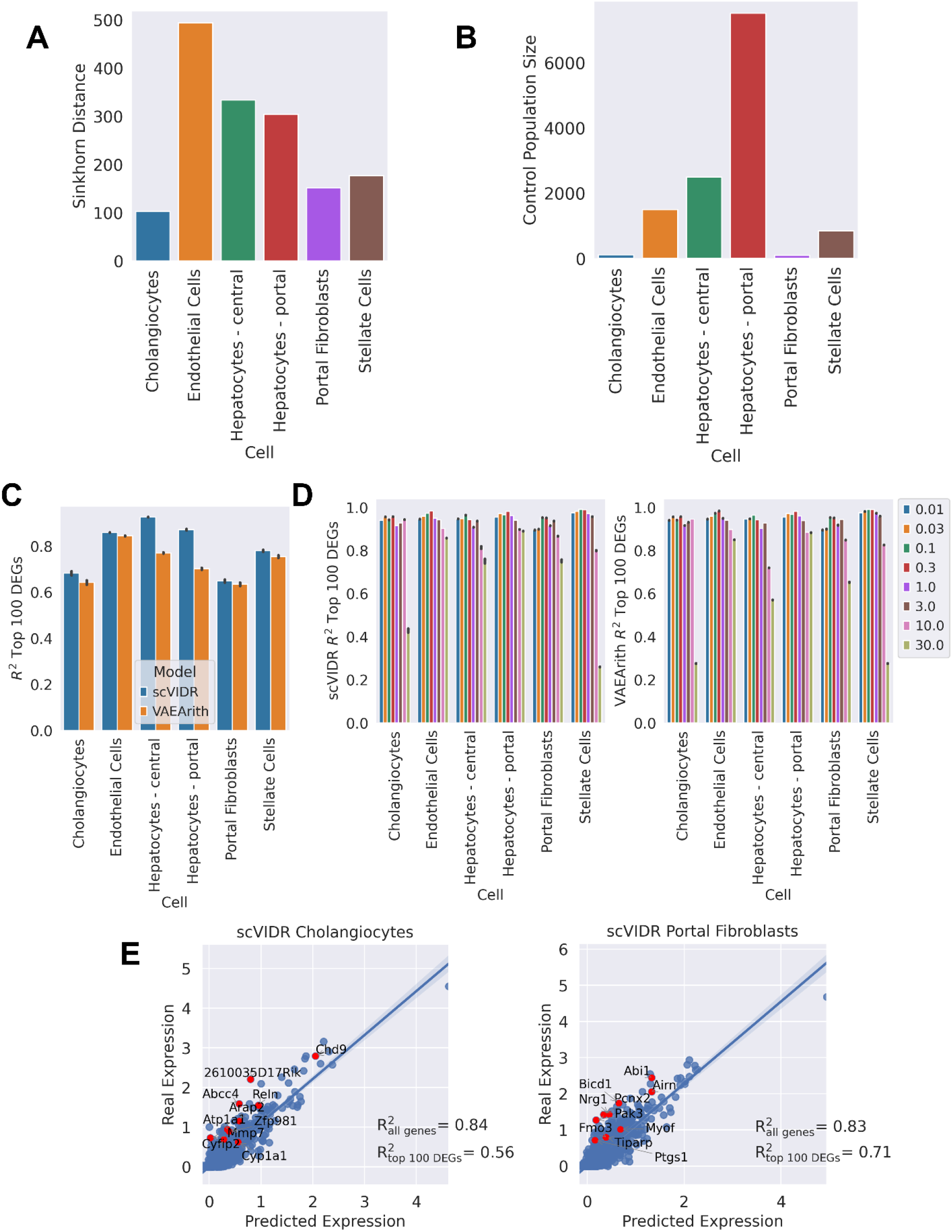
The relationship between statistical distance between control and treated distributions on the latent space and overall model performance in mouse liver treated with TCDD. **A)** Sinkhorn distance between the latent distributions of the control and 30 *μ*g/kg doses of TCDD of each cell type on the latent space. **B)** Bar plot of the control group cell population size for each cell type. **C)** Bar plot of mean gene *R*^2^ for each individual cell type when predicting only the 30 *μ*g/kg dose of TCDD. **D)** Bar plot of mean *R*^2^ for each individual cell type when predicting across the entire TCDD dose-response experiment. **E)** scVIDR prediction versus real expression regression plot of cholangiocytes and stellate cell from mice administered with a 30 *μ*g/kg dose of TCDD. Each point represents the mean expression of a gene. The top 10 differentially expressed genes are represented with red points.

**Supplementary Figure 6.**
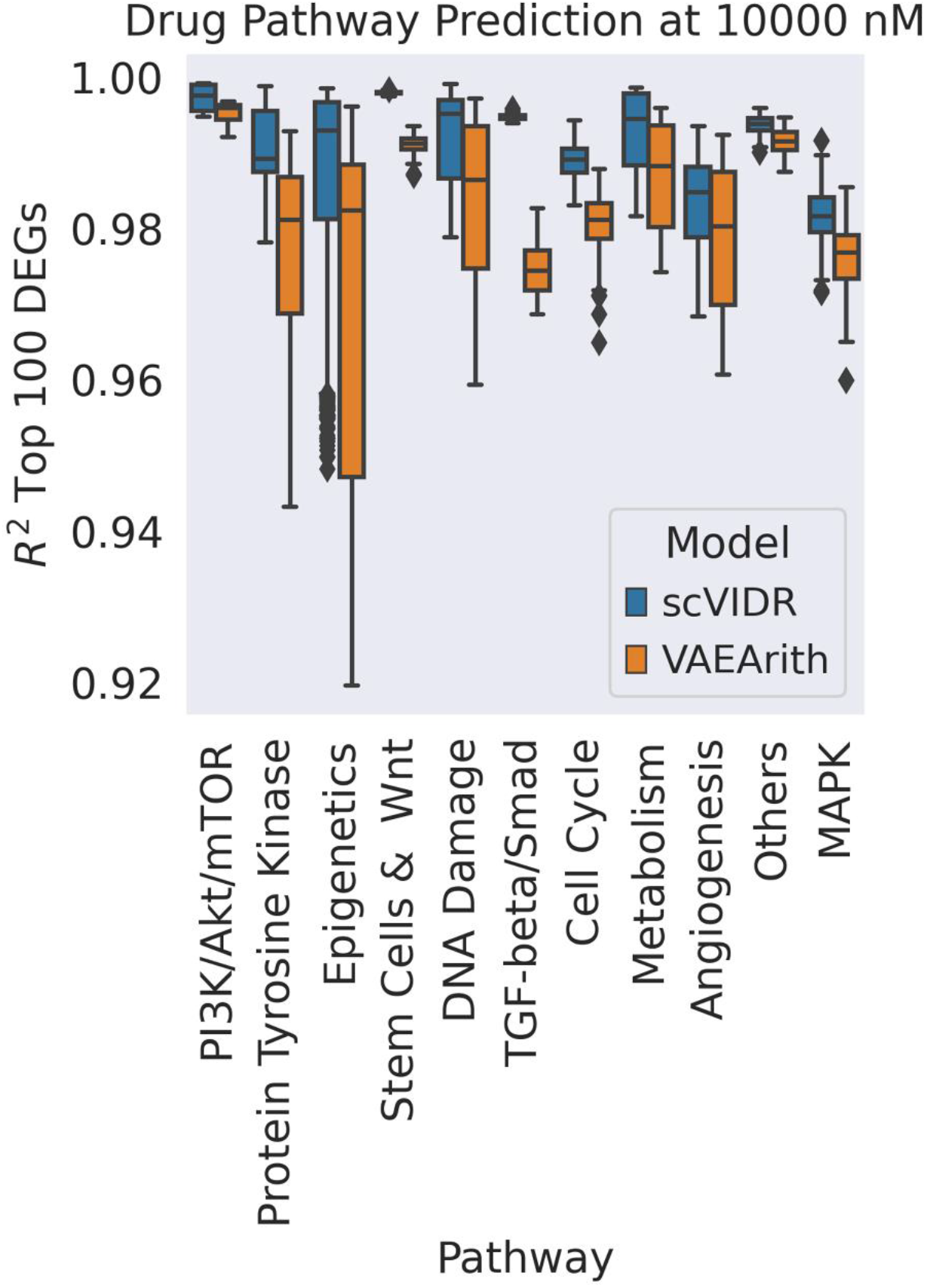
Overall drug pathway performances at the highest administered dose. **A)** A boxplot of the mean gene *R*^2^ across all drug pathways in the test dataset at a dose of 10,000 nM.

## Notes

### Competing Interest Statement

The authors have declared no competing interest.

